# Researcher effects on the biological structure and edaphic conditions of field sites and implications for management

**DOI:** 10.1101/2023.06.23.546286

**Authors:** Shelby A Rinehart, Jacob M Dybiec, Parker Richardson, Janet B Walker, James D Peabody, Julia A Cherry

**Author notes:** Authors contributed equally (Shelby A Rinehart, Jacob M Dybiec, Parker Richardson, Janet B Walker). **Corresponding Author:** Julia A Cherry Box 870344; The University of Alabama; Tuscaloosa, AL 35487-0344. **Contact Author:** Shelby A Rinehart Box 870344; The University of Alabama; Tuscaloosa, AL 35487-0344. **Open Research Statement:** Upon acceptance, all data will be provided through Dryad.

## Abstract

Field studies are necessary for understanding natural processes, but they can disturb the environment. Despite researchers acknowledging these effects, no studies have empirically tested the direct (e.g., harvesting plants) and indirect effects (i.e., trampling) of researcher activities on biological structure and edaphic conditions. We leveraged field studies in Alabama and California to monitor the recovery of tidal marshes following research activities. Researcher effects on animals, plants, and sediment conditions remained prevalent almost one year after the disturbance ended. For instance, trampled plots had 14-97% lower plant cover than undisturbed plots after >10 months of recovery. Researcher effects also impacted plant composition, leading to increased subordinate species abundance. We encourage field researchers to adopt strategies that reduce their scientific footprints, including reducing field visits, limiting field team size, and considering ways to limit potential environmental impacts during study design.

## Introduction

Experimental and observational field studies are essential tools for developing ecological theories that describe the processes governing ecosystem structure and functions. For instance, field studies have helped develop island biogeography theory (Simberloff & Wilson 1969; Simberloff 1974), the resource ratio hypothesis (e.g. The Park Grass experiment; Tilman 1982; Silvertown et al. 2006), the keystone species concept (Paine 1966), theories of species interactions (Connell 1961; Estes & Palmisano 1974), and models of ecosystem disturbance (e.g., Hubbard Brook Ecosystem Study; Holmes and Likens 2016). Despite field studies being critical tools to advance scientific understanding, as a community of field researchers, we would be remiss if we did not recognize that conducting field research can disturb and degrade important habitats and the species that inhabit them (Bryzek et al. 2022). Here, we evaluate the effects of researchers following a series of manipulative field experiments to better understand the impacts of field research and develop practices to mitigate these impacts.

Field researchers can disturb ecosystems through direct (i.e., scientific activities) and indirect (e.g., unintentional activities) pathways. The direct effects of researchers come in two key forms. First, sample collection can disturb various components of the environment. For instance, collecting and interacting with wild animals can have adverse effects on their populations and behaviors (Ikin 2011; China et al. 2021; Lewis et al. 2022). Additionally, harvesting plants can defoliate habitat, alter plant community composition, and impact nutrient processing rates (Hille et al. 2018; Jabłońska et al. 2021). Soil coring and soil pits may also adversely affect environmental processes. Specifically, coring sediments and soils may be analogous to animal burrowing, increasing sediment homogenization and oxygenation (Bertness 1985; Bortolus & Iribarne 1999; Kristensen & Alongi 2006; Rinehart et al. 2023a).

Second, planned manipulations of environmental attributes during field experiments, while designed to test specific hypotheses, may create legacy effects (intentional and unintentional) on local biological structure and ecosystem functions that can last beyond the life of the study. For example, David Simberloff and E.O. Wilson conducted a study wherein they fumigated entire mangrove islands along the Florida Keys with methyl bromide to remove all arthropod fauna so they could monitor island re-colonization (Simberloff & Wilson 1969; Simberloff et al. 1970). This study led to rapid mangrove damage (i.e., 5-100% of leaves were burnt; Wilson & Simberloff 1969), a decline in arthropod abundance that lasted over 250 days (Simberloff & Wilson 1969) and a shift in arthropod community composition that lasted at least two years (Simberloff et al. 1970). Similarly, Deegan et al. (2012) enriched ∼30,000 m^2^ of tidal marsh in the Plum Island Estuary in Massachusetts with nitrogen and phosphorus over a nine-year period to test the effects of eutrophication on coastal ecosystems. Their manipulation resulted in decreased belowground productivity, higher rates of creek-bank collapse, and the conversion of vegetated marsh into unvegetated mud flats, all of which can degrade tidal marsh ecosystem services (e.g., carbon sequestration and nitrogen removal; Craft et al. 2009; Deegan et al. 2012; Hinshaw et al. 2017). Given the potential extent and longevity of researcher direct effects, it is important to consider the consequences of field manipulations for biological structure and ecosystem functions.

Researcher indirect effects occur when researchers impact the environment through activities such as trampling. Human trampling can affect the physiology and morphology of vegetation (Kuss & Graefe 1985; Goldman Martone & Wasson 2008), subterranean fauna (Chappell et al. 1971) and edaphic conditions (Ros et al. 2004; Pescott & Stewart 2014). Additionally, the impacts of human trampling on vegetation depends on the specific traits (e.g., life forms) of the plant populations and communities (Pescott & Stewart 2014). For instance, habitats dominated by grasses (e.g., *Festuca* spp.) have been shown to be less susceptible to the impacts of human trampling than habitats dominated by deciduous shrubs (e.g., *Symphoricarpos* spp.) and herbaceous perennial plants (e.g., *Clintonia* spp.; Cole 1987). Thus, broadly evaluating the impacts of researcher trampling on plants requires considering the variation in plant traits observed across ecological communities.

Despite the clear pathways by which researcher activities (direct and indirect) may impact the environment, to our knowledge, there have been no attempts to quantify the direct and indirect effects of researcher disturbance in ecological systems. This gap is surprising given that 1) various field researcher communities (e.g., hydrothermal vent researchers) have implemented codes of conduct to limit researcher impacts (Devey et al. 2007; Godet et al. 2011; Van Dover 2012; Van Dover et al. 2012; Bryzek et al. 2022) and 2) similar human activities (e.g., recreational habitat trampling) have well-documented negative effects across habitats (Cole 1987, 1995; Leung & Marion 2000; Pescott & Stewart 2014). Thus, empirically testing how researcher effects (direct and indirect) influence the environment will provide a critical first step in developing a framework for implementing ethical and sustainable field practices.

Here, we evaluated the direct and indirect effects of researchers in natural and constructed tidal marshes along the Alabama and southern California coastlines. Specifically, we monitored the recovery of animal populations, plant communities, and sediment conditions for approximately one year in areas that were previously 1) manipulated and sampled (e.g., defoliated) and 2) impacted by researcher trampling. We predicted that researcher indirect effects would be greater than direct effects in all ecosystems. Additionally, we anticipated that researcher effects would be more pronounced in constructed marshes than in natural marshes because constructed systems lack the material and informational legacies necessary for rapid recovery following disturbance (Johnstone et al. 2016). Understanding how researcher activities impact ecosystems with different natural histories (e.g., natural versus constructed) in distinct North American ecoregions will help inform predictions of when and where researchers have detrimental effects on the environments they study and hopefully lead to the development and implementation of ethical practices to mitigate those impacts.

## Methods

### Conceptual approach

In 2018 and 2021, we completed caging studies manipulating the presence and absence of burrowing crab communities in tidal marshes in two regions—southern California (CA) and Alabama (AL), respectively (Walker et al. 2021a, 2021b; Appendix S1). Both studies involved manipulating crab population densities using cages (i.e., high crab and low crab burrow densities), harvesting aboveground plant biomass (CA: 0.7 x 0.7m, length x width; AL: 0.5 x 0.5m), coring sediments (CA: 50 x 50 x 27cm, length x width x depth; AL: 18 x 10cm, diameter x depth), and severing plant rhizomes (CA: 30 cm depth; AL: 15 cm depth; Appendix S1). Maintaining and sampling these experimental manipulations resulted in habitat trampling. Following our direct (i.e., manipulations and sampling) and indirect (i.e., trampling) effects on tidal marshes in AL and CA, we monitored the recovery of crab populations, plant community structure, and edaphic conditions (AL only) through the following growing season in each marsh site. To minimize the environmental impacts of our continued monitoring we 1) stayed on established trails, 2) reduced team sizes (1-2 researchers per visit), and 3) decreased our site visits to every other month.

### Site descriptions

In both regions, we used two tidal marsh sites. Each region had one natural tidal marsh and one constructed tidal marsh (AL: 34 years old; CA: ∼17 years old).

#### Alabama

Our natural site was Fowl River Natural Marsh (AL-NAT; 30°22’02.5’’ N, 88°09’37.2’’ W) and our constructed site was Fowl River Constructed Marsh 1 (AL-CON; 30°22’02.3’’ N, 88°09’06.8’’ W). AL-CON was initially constructed in 1987 by harvesting pine savanna habitat and excavating topsoil down to a clay layer that intercepted the water table and was 0.27m above mean sea level (NAVD88 datum). The site was then planted with needlerush (*Juncus roemerianus*) and smooth cordgrass (*Spartina alterniflora*) in 1988 (Vittor et al. 1987). Today, both AL sites are almost entirely dominated by needlerush (Ledford et al. 2021; Tatariw et al. 2021; Rinehart et al. 2023a), although the subordinate species, *Distichlis spicata*, is also common at AL-NAT. Burrowing crabs, fiddler crabs (i.e., *Minuca longisignalis*, *Minuca minax*, *Leptuca panacea*, and *Leptuca spinicarpa*) and marsh crabs (*Sesarma reticulatum*) are common and have known effects on needlerush productivity and sediment conditions (Rinehart et al. 2023a).

#### Southern California

Our natural site was Kendall-Frost Marsh (CA-NAT; 32°47041.0’’ N, 117°13046.4’’ W) and our constructed site was San Dieguito Lagoon (CA-CON; 32°58047.0’’ N, 117°14043.6’’ W). CA-CON was initially constructed in 2006 and planted with Pacific cordgrass (*Spartina foliosa*) and pickleweed (*Sarcocornia pacifica*) in 2008, with subsequent grading and planting activities occurring from 2009-2022 (Beheshti et al. 2022). At both sites, Pacific cordgrass dominates low-elevation habitat, pickleweed dominates high-elevation habitat, and mixtures of these species occur at intermediate elevations (Walker et al. 2021b, 2022). Subordinate plants are more common at CA-NAT and include *Salicornia bigelovii, Jaumea carnosa, Batis maritima,* and *Triglochin maritima*. Burrowing crabs, including lined shore crabs (*Pachygrapsus crassipes*) and Mexican fiddler crabs (*Leptuca crenulata*), are abundant at CA-NAT and CA-CON and can influence plant community structure and edaphic conditions (Walker et al. 2021a, 2021b).

### Experimental design

The study included four treatments: High Crab, Low Crab, Trampled, and Control (n = 5 treatment^-1^) at constructed and natural marshes in both regions. The High Crab and Low Crab treatments were established in the footprints of the high-crab density and low-crab density cages, respectively, that were part of the original burrowing crab manipulative studies conducted at each site (Appendix S1). Thus, these plots had their crab communities modified, aboveground biomass harvested, rhizomes severed, and sediments cored as part of prior experiments (Appendix S1). Trampled plots were established in areas where researchers frequently walked to maintain the original caging studies. Controls were placed in nearby habitat that was previously undisturbed by researcher activities. Plots sizes were 0.7 x 0.7m (length x width) in CA and 0.5 x 0.5m in AL.

Starting in February (AL: 2021; CA: 2019), we monitored crab burrow density, plant cover, and the mean height and stem density (AL: n ≤ 5 stems plot^-1^; CA: n ≤ 10 stems plot^-1^) of the dominant plant species (AL: needlerush; CA: Pacific cordgrass) in each marsh every-other month until September (CA) and October (AL). Additionally, in October in AL marshes, we collected one, 10-cm-deep sediment core using a Russian Peat corer (i.d. = 5cm) that we sub-sectioned in the field at 2.5cm intervals. Subsections were oven-dried at 60°C to a constant mass to obtain bulk density, and once dried, samples were ground with a mortar and pestle before being ashed in a muffle furnace (6h at 550°C) to estimate sediment organic matter (SOM) via loss on ignition. We also collected one, 10-cm-deep sediment core using a t-corer (i.d. = 7.9cm) that we sub-sectioned in the field at 5 cm intervals to assess belowground biomass at the AL sites. Subsections were rinsed to remove sediment attached to belowground biomass, which was then dried at 60°C to a constant mass (Conner & Cherry 2015).

### Data analysis

We used separate two-factor ANOVAs for each region (i.e., AL and CA) to evaluate the effects of researcher impacts (i.e., experimental manipulations and trampling) on tidal wetland burrowing crab populations, plant communities, and edaphic conditions in each region. However, we were unable to evaluate crab burrow density at AL-CON. Because of this, we modified our analysis and ran a one-factor ANOVA for burrowing crab density between researcher impact treatments at AL-NAT. All models included treatment and site (i.e., NAT, CON) as fixed factors. We ran separate models for each region because the studies were conducted in different years and have distinct plant communities. If model assumptions were violated, we transformed the data. We used Tukey’s HSD tests (α = 0.05) when significant variables were detected. We used measurements taken at the end of each region’s growing season (AL = October; CA = September) in all ANOVAs, thus time was not a factor in any of our models. All temporal data is available in Rinehart et al. (2023b).

We also used separate two-factor ANOVAs for each AL site (i.e., NAT and CON) to assess the effects of the experimental manipulations on belowground biomass, sediment bulk density, and SOM at the end of the growing season (October) in AL. We ran separate models for each AL site because we knew that AL-CON had less belowground plant biomass, higher sediment bulk density, and lower SOM content than AL-NAT (Ledford et al. 2021; Tatariw et al. 2021; Rinehart et al. 2023a). Thus, models included treatment and sediment depth as fixed factors. Tukey’s HSD tests (α = 0.05) were used as needed. All models were conducted with the “aov” function in R software v. 4.0.2 (Kassambara 2020, R-Core-Team 2020).

We compared the effects of experimental manipulations and trampling across regions by estimating the log response ratio (LRR; mean ± 1var) effect size in OpenMEE (Wallace et al. 2017). LRRs were calculated using the High Crab, Low Crab, and Trampled treatments as experimental groups and the Control treatment as the control group (Hedges et al. 1999; Lajeunesse 2011). We calculated LRRs for all crab population, plant community, and edaphic condition metrics. Specifically, with plant community response variables, we calculated LRRs for the stem height and stem density of the dominant plant species in each region (i.e., needlerush in AL and Pacific cordgrass in CA). Negative LRRs suggest researcher effects decrease the response variable relative to undisturbed controls, while positive LRRs suggest researcher effects increase the response variable relative to undisturbed controls. LRRs are presented as a forest plot. We generalized researcher impacts between sites by qualitatively comparing how the LRRs for each response variable deviated from zero.

## Results

### Alabama

#### Crab populations

Crab burrow density data were only available for the first two (February and April) of the five months sampled, thus we removed AL-CON from the analysis and ran a one-factor ANOVA with treatment as fixed factor for crab burrow density at AL-NAT. We were also unable to collect burrow density data from one replicate of each treatment at AL-NAT in October due to tidal inundation, thus all treatments had a n = 4 for this analysis. Overall, researchers had no effect on crab burrow density at AL-NAT (F *=* 2.23, df = 3, p = 0.138). However, researcher direct effect plots (High Crab, Low Crab) tended to have higher crab burrow densities than Trampled and Control plots. Additionally, LRRs suggest that researcher direct (i.e., manipulations and sampling) and indirect effects (i.e., trampling) have a slight positive effect on crab burrow density (Figure 1).

**Figure 1.**
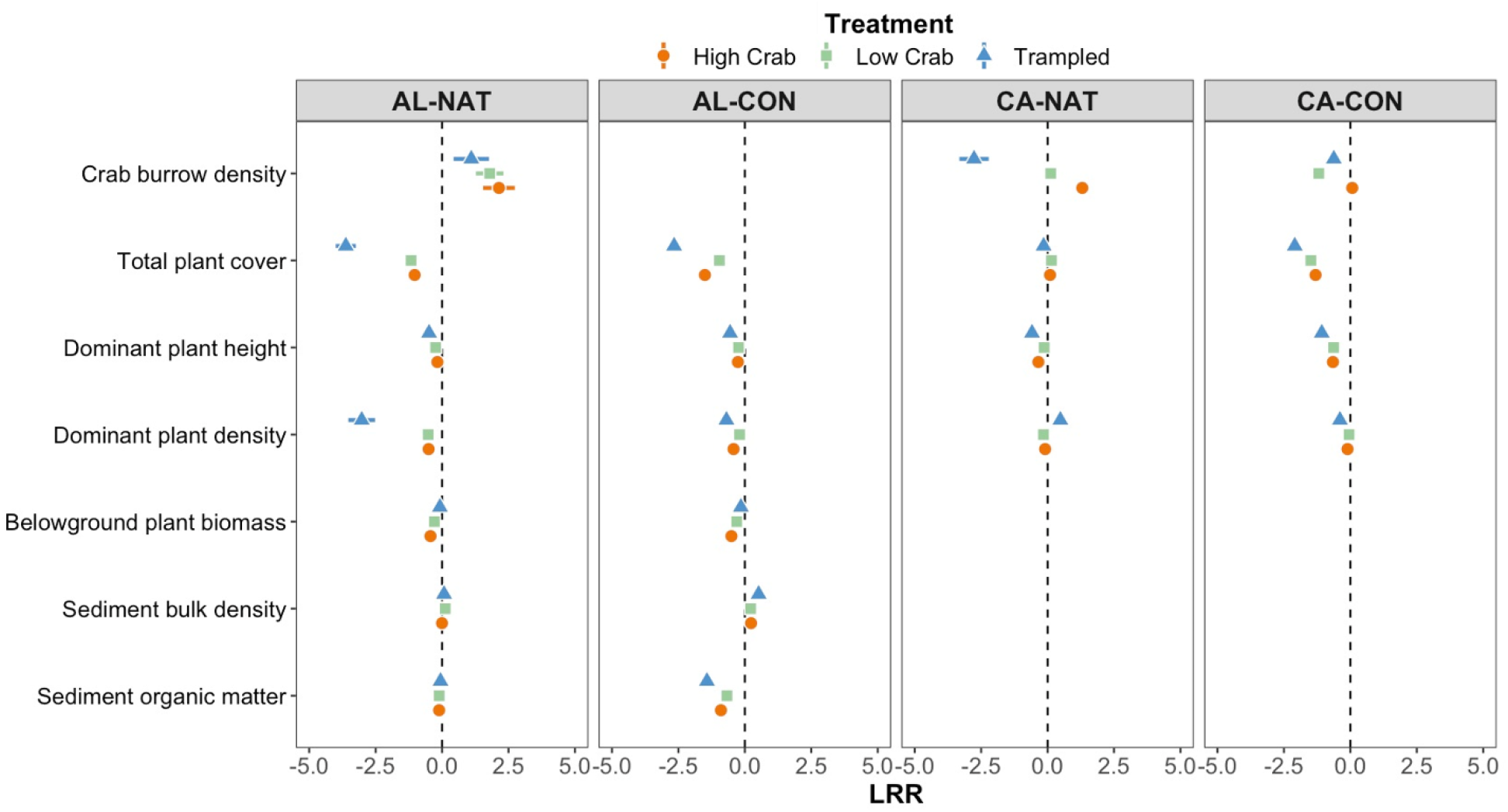
Log response ratio (LRR) effect size calculations (mean ± 1 var) for the impact of researcher effects treatments (i.e., High Crab, Low Crab, and Trampled) on burrowing crab populations, plant community traits, and sediment properties in four tidal marshes. High Crab treatments are denoted by orange circles, Low Crab treatments are denoted by green squares, and Trampled treatments are denoted by blue triangles. AL-NAT and AL-CON are the natural and constructed tidal marshes, respectively, in Alabama. CA-NAT and CA-CON are the natural and constructed marshes, respectively, in southern California.

#### Plant communities

There was an interaction between Marsh and Treatment on total plant cover (F = 4.44, df = 3, p = 0.010; Figures 1-3; Appendix S2: Table S1). At AL-NAT and AL-CON, all researcher impact plots (High Crab, Low Crab, and Trampled) had lower total plant cover than Control plots. Additionally, Trampled plots had lower plant cover than High Crab and Low Crab plots at AL-NAT and Trampled and High Crab plots had lower plant cover than Low Crab plots at AL-CON (Appendix S3: Figure S1). LRRs suggest that researcher direct and indirect effects have a large negative effect on total plant cover (Figure 1).

**Figure 2.**
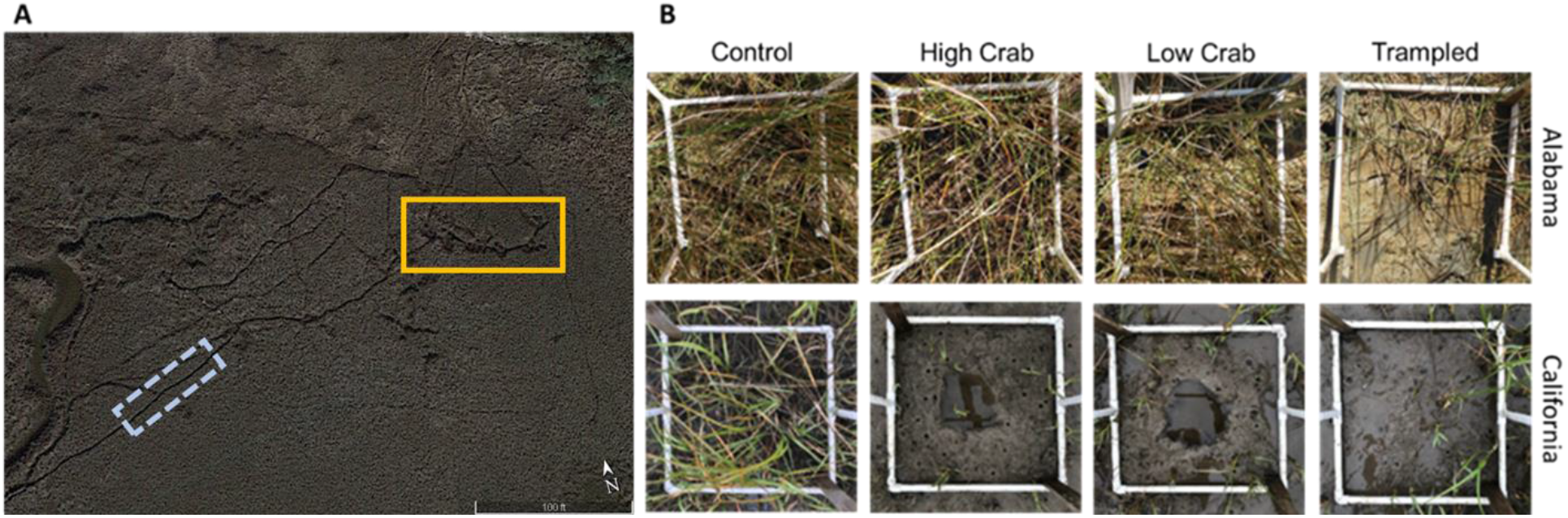
A) Google Earth image of the Alabama natural marsh (AL-NAT) from December 2022. The blue rectangle (dashed line) denotes researcher trails caused by trampling. The yellow rectangle (solid line) denotes the footprint of the burrowing crab caging study deployed from April – December 2021 (Appendix S1). B) Images of each treatment (i.e., Control, High Crab, Low Crab, and Trampled) at the constructed tidal marshes (i.e., AL-CON, CA-CON) in each region. Images in the top row represent plots from the Alabama constructed marsh (AL-CON). Images in the bottom row represent plots in the California constructed marsh (CA-CON). Images are from October 2022 in Alabama and June 2019 in California, ten months and eight months, respectively, after the researcher effects occurred.

**Figure 3.**
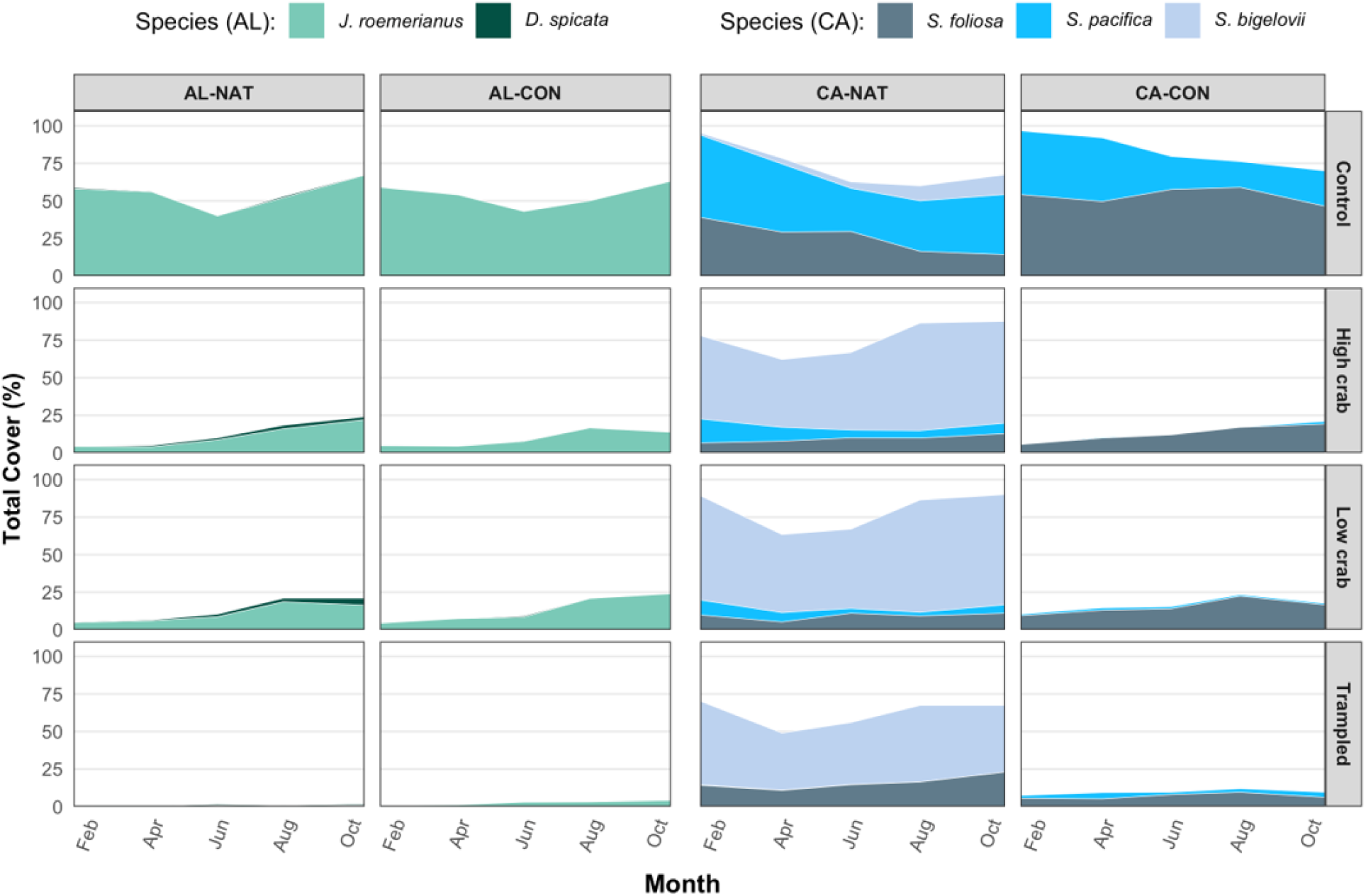
The mean total cover (%) of plants in each treatment (z-axis; Control, High Crab, Low Crab, Trampled) at each natural (NAT) and constructed (CON) tidal marsh in Alabama (AL) and California (CA) over the growing season (February – October). Each color represents a distinct plant species. For AL-NAT and AL-CON, light green represents the relative percent cover of *Juncus roemerianus* (needlerush) and dark green represents the relative percent cover of *Distichlis spicata*. For CA-NAT and CA-CON, dark blue represents the relative percent cover of *Spartina foliosa* (Pacific cordgrass), aqua represents the relative percent cover of *Sarcocornia pacifica* (pickleweed), and light blue represents the relative percent cover of *Salicornia bigelovii.*

Marsh and Treatment interacted to affect needlerush stem density (F = 5.22, df = 3, p = 0.005; Appendix S2: Table S1). At AL-CON, needlerush stem density was similar in High Crab, Low Crab, and Control plots. However, Trampled plots had 24-50% fewer needlerush stems than High Crab, Low Crab, and Control plots (Figure 3A). At AL-NAT, High Crab, Low Crab and Trampled plots had 40-95% fewer needlerush stems than Control plots (Figure 3A). High and Low Crab plots also had higher needlerush stem densities than Trampled plots. LRRs suggest that researcher effects have negative effects on needlerush stem density, but researcher indirect effects tend to be more detrimental than researcher direct effects (Figure 1).

Treatment affected needlerush stem height (F = 24.16, df = 3, p < 0.001; Figure 3). In both marshes, stems in High Crab, Low Crab, and Trampled plots were shorter than stems in Control plots (Figure 3b). Needlerush stems in High Crab and Low Crab plots were also taller than stems in Trampled plots (Figure 3b). LRRs suggest that researcher direct and indirect effects have a slight negative effect on mean needlerush stem height (Figure 1).

At AL-NAT, there was no significant effect of Treatment or Depth on plant belowground biomass. However, there was an interaction between Treatment and Depth on plant belowground biomass at AL-CON (F = 3.25, df = 3, p = 0.034; Appendix S2: Table S2; Appendix S3: Figure S2). Trampled plots tended to have higher belowground biomass at 0-5 cm depth than High Crab, Low Crab, and Control plots; but this effect diminished at depths of 5-10 cm (Appendix S3: Figure S2). Across all depths, LRRs suggest that researcher indirect effects did not impact plant belowground biomass, but researcher direct effects have a slight negative effect on plant belowground biomass in both marshes (Figure 1).

#### Sediment characteristics

Bulk density increased with depth at both marshes (AL-NAT: F = 7.75, df = 3, p < 0.001; AL-CON: F = 3.60, df = 3, p = 0.018; Figure 5; Appendix S2: Table 2). Researcher effects (Treatment) did not impact sediment bulk density at AL-NAT (F = 0.75, df = 3, p = 0.527; Appendix S2: Table S2), but they did at AL-CON (F = 4.66, df = 3, p = 0.005; Appendix S2: Table S2). Specifically, at AL-CON, sediment bulk density was 68% higher (averaged across all depths) in Trampled plots than Control plots (Figure 5). LRRs indicate that researcher effects did not affect bulk density at AL-NAT, but they had slight negative effects on bulk density at AL-CON (Figure 1).

**Figure 4.**
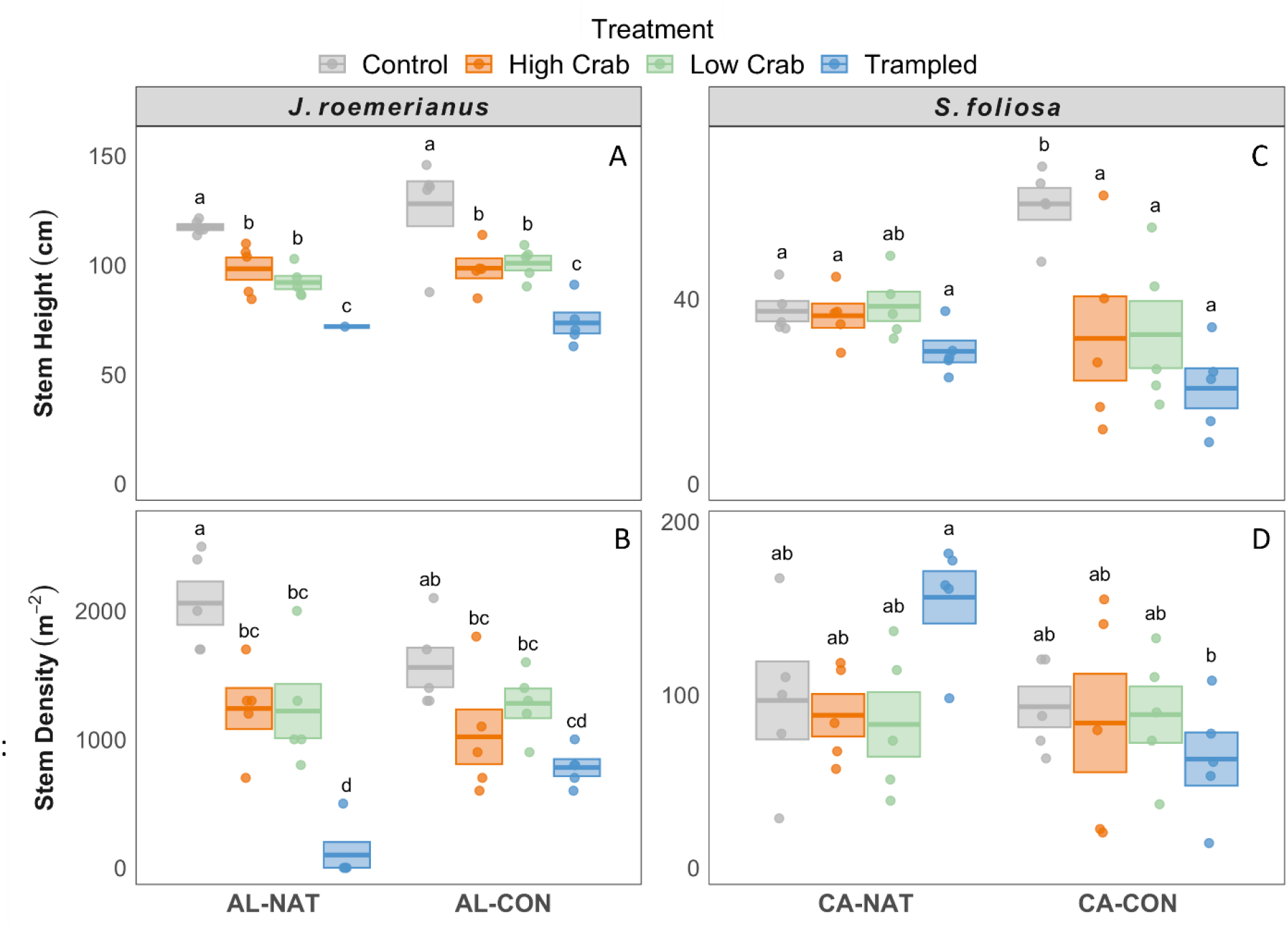
A) Mean stem height and B) stem density of *Juncus roemerianus* (needlerush) at the end of the growing season in all treatments (Control, High Crab, Low Crab, and Trampled) at the Alabama (AL) natural marsh and constructed marsh (AL-NAT and AL-CON, respectively). C) Mean stem height and B) stem density *Spartina foliosa* (Pacific cordgrass) at the end of the growing season in all treatments (Control, High Crab, Low Crab, and Trampled) at the California (CA) natural marsh and constructed marsh (CA-NAT and CA-CON, respectively). Mean stem height is based on the mean of five needlerush stems in AL and ten Pacific cordgrass stems in CA. Lines inside boxes are mean values, box limits represent ± 1 SE. Points represent raw data. Letters represent significant interactions between site and treatment (Tukey HSD test; α = 0.05).

**Figure 5.**
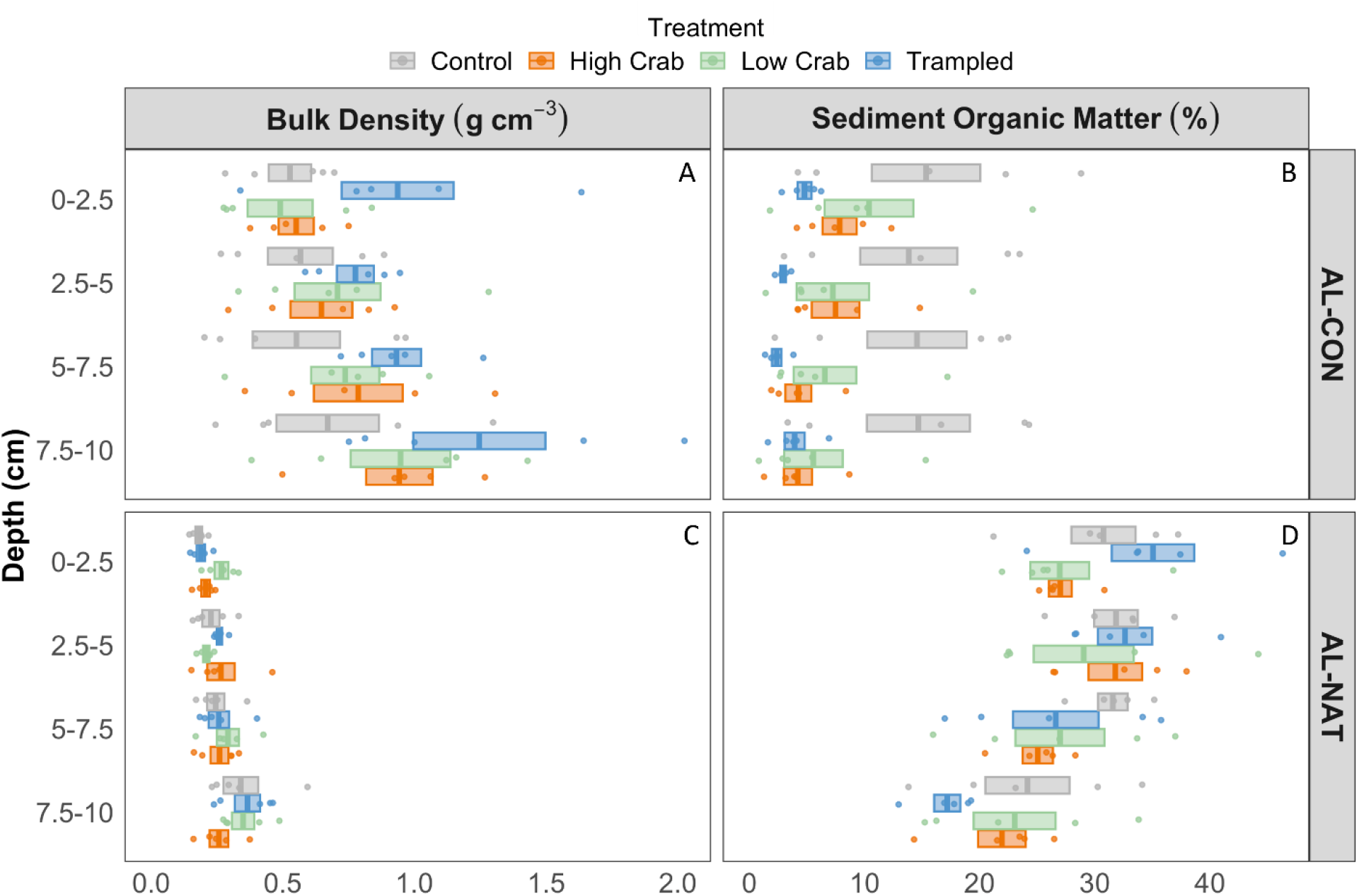
A) Sediment bulk density and B) sediment organic matter (SOM) by depth in all treatments (Control, High Crab, Low Crab, Trampling) at the constructed (CON) marsh in Alabama. C) Sediment bulk density and D) sediment organic matter (SOM) by depth in all treatments (Control, High Crab, Low Crab, Trampling) at the natural (NAT) marsh in Alabama. Lines inside boxes are mean values, box limits represent ± 1 SE. Points represent raw data.

At AL-NAT, SOM decreased with depth (F = 9.48, df = 3, p < 0.001; Figure 5; Appendix S2: Table S2). Researcher effects did not impact SOM at AL-NAT (F = 1.11, df = 3, p = 0.350; Appendix S2: Table S2). Depth did not affect SOM at AL-CON (F = 0.74, df = 3, p = 0.531; Appendix S2: Table S2). However, researcher direct and indirect effects decreased SOM at AL-CON by 49-76% relative to undisturbed Control plots (F = 11.40, df = 3, p < 0.001; Figure 5; Appendix S2: Table S2). LRRs indicate there were no researcher effects on SOM at AL-NAT, but there were substantial negative effects of researchers on SOM at AL-CON (Figure 1).

### California

#### Crab populations

Crab burrow density was impacted by Marsh (F = 7.42, df = 1 p = 0.011), Treatment (F = 3.71, df = 3, p = 0.023), and their interaction (F = 2.99, df = 3, p = 0.048). CA-CON had 49% more crab burrows across all treatments than CA-NAT. Treatments had distinct impacts on crab burrow density at each Marsh. Specifically, at CA-CON, there was no effect of Treatment on crab burrow density, while at CA-NAT, Trampled plots had fewer crab burrows than Control, High Crab, and Low Crab plots. These findings were supported by our LRRs, which suggest that researcher impacts have small negative effects on crab burrow density at CA-CON and have variable effects on crab burrow density at CA-NAT. Specifically, trampling decreased burrow density, High Crab manipulations increased burrow density, and Low Crab manipulations had no effect on burrow density (Figure 1).

#### Plant communities

Marsh (F = 248.3, df = 1, p < 0.001), Treatment (F = 25.1, df = 3, p < 0.001), and their interaction (F = 25.44, df = 3, p < 0.001) influenced total plant cover (Appendix S2: Table S1). At CA-NAT, plant cover was similar across all treatments, suggesting that plant cover had recovered to control conditions within one year of researcher impacts (Figure 1, 3; Appendix S2: Table S1; Appendix S3: Figure S1). However, High Crab, Low Crab, and Trampled plots at CA-CON had 73-88% less total plant cover than Control plots after a year of recovery (Figures 1-3; Appendix S2: Table S1; Appendix S3: Figure S1). LRRs suggest that researcher direct and indirect effects had no impact on total plant cover at CA-NAT but decreased total plant cover at CA-CON (Figure 1).

CA-NAT had 29% more Pacific cordgrass stems across all treatments than CA-CON (F = 3.42, df = 1, p = 0.070; Figure 4). There was also an interaction between Marsh and Treatment on Pacific cordgrass stem density (F = 3.23, df = 3, p = 0.040; Appendix S2: Table S1). This interaction was mediated by different effects of trampling on Pacific cordgrass stem density at each marsh. Pacific cordgrass stem density was 32% lower in Trampled plots than Control plots at CA-CON, while it was 62% higher in Trampled plots than Control plots at CA-NAT (Figure 4). The opposing effect of researcher trampling on cordgrass stem density between CA-CON and CA-NAT is further supported by our LRRs, which suggest that researcher direct effects have minimal impacts on Pacific cordgrass stem density at both marshes. However, researcher indirect effects have a slight negative effect on Pacific cordgrass stem density at CA-CON and a slight positive effect on Pacific cordgrass stem density at CA-NAT (Figure 1).

Treatment also affected Pacific cordgrass stem height (F = 6.63, df = 3, p = 0.001; Figure 4; Appendix S2: Table S1). There was no effect of Marsh or Marsh by Treatment interactions on Pacific cordgrass stem height (Appendix S2: Table S1). Pacific cordgrass stems in High Crab, Low Crab, and Trampled plots were 47-66% shorter than stems in Control plots at CA-CON and were 12-44% shorter than stems in Control plots at CA-NAT (Figure 3). LRRs suggest that researcher activities have negative effects on Pacific cordgrass stem heights, with these effects being greater at CA-CON than CA-NAT (Figure 1).

## Discussion

We show that researcher direct effects (i.e., sampling and manipulating) and indirect effects (i.e., trampling) have pervasive impacts on crab burrow density, plant cover and traits, and edaphic conditions in Alabama and southern California tidal marshes. In all marsh sites, researcher indirect effects had greater impacts on plant communities and sediment conditions than researcher direct effects (Figure 1). Additionally, researcher effects (direct and indirect) tended to be greater in Alabama than in southern California tidal marshes, possibly due to differences in the functional traits of dominant plants in the communities of these regions. Research impacts were also greater in constructed tidal marshes than natural tidal marshes across both regions, suggesting that constructed tidal marshes are less resilient to disturbance than natural tidal marshes. Our study highlights the detrimental effects that research-related activities can have on the biological structure and edaphic conditions of field sites and provides unique insights that can help researchers minimize the scientific footprint of their research programs.

Researcher indirect effects associated with trampling had a greater impact on tidal marsh plant community structure and edaphic conditions than researcher direct effects (e.g., manipulated plots; High Crab and Low Crab plots). The profound impacts of researcher trampling on tidal marsh structure are not surprising for two reasons. First, studies with other large mammals (e.g. reindeer, sheep, cattle) have repeatedly found that mammal trampling has greater impacts on the abundance and composition of plant communities than defoliation from mammal grazing (Sørensen et al. 2009; Lezama & Paruelo 2016; Narantsetseg et al. 2018). For example, reindeer trampling in a sub-arctic grassland decreased the cover of moss (*Pleurozium schreberi*) and sedges (*Carex vaginata*), while reindeer defoliation did not affect plant cover (Sørensen et al. 2009). Thus, the indirect effects of mammals, broadly, on their environments may be greater than their direct effects.

Second, human vegetation trampling associated with recreational activities has well-documented, adverse effects on plant community structure and sediment conditions (Goldman Martone & Wasson 2008; Pescott & Stewart 2014). Cole (1987) found that plant cover in grasslands and forests tends to decline exponentially with increasing human trampling intensity (i.e., number of passes per year). In fact, 43% of plant cover was lost after only 25-40 passes per year (Cole 1987). This loss in plant cover is similar to the trampling effects observed in our Alabama sites, where approximately 33 passes (per year) led to a 39-43% decline in plant cover in Trampled plots relative to Control plots (Figure 3). Thus, studies of recreational human trampling may provide valuable insights into how researcher trampling impacts plant structure and edaphic conditions at field sites.

While indirect trampling effects were the main pathway by which researchers affected field sites, researcher direct effects did have legacy effects on crab burrow density and belowground plant biomass. For instance, at CA-NAT and AL-NAT, High Crab plots still had 269% and 750%, respectively, more crab burrows than Control plots 10-12 months after the final crab manipulation occurred (Appendix S3: Figures S3-S4). Elevated crab burrow densities can affect plant community composition, sediment homogenization, sediment oxygenation, organic matter distribution, and decomposition (Bertness 1985; Bortolus & Iribarne 1999; Gribsholt & Kristensen 2002; Kristensen & Alongi 2006; Walker et al. 2021a, 2021b; Rinehart et al. 2023a). Combined, these results suggest that researcher manipulations of animal populations, such as burrowing crabs, may have long-term cascading effects on the biological structure and ecosystem functions of field sites. While the direct effects of field manipulations are often intentional (e.g., shifting crab burrow densities), the spatial and temporal scales of these impacts should be considered when designing field studies.

Plant functional traits (e.g., life-form, growth-form) frequently affect the resistance and resilience of plant communities to human disturbance and trampling (see Cole 1987, 1995; Pescott & Stewart 2014). In fact, plant functional traits are better predictors of trampling effects than the intensity of the trampling, since low intensity trampling can be as damaging as high intensity trampling in some plant communities (Pescott & Stewart 2014). Our study also supports this observation, since natural tidal marshes dominated by Pacific cordgrass had greater resiliency to researcher effects than marshes dominated by needlerush (Figure 1). For instance, after one growing season, Pacific cordgrass stem density was 62% higher in Trampled plots than Control plots at CA-NAT, while needlerush stem density was still 95% lower in Trampled plots than Control plots at AL-NAT (Figure 4). Pacific cordgrass is likely more resilient to researcher effects (direct and indirect) than needlerush because it is more stress tolerant and has faster rates of clonal spread (Touchette et al. 2009; Jones et al. 2016), and as such, our Alabama and southern California sites may require different approaches to mitigate researcher impacts and facilitate recovery. Thus, it is important that researchers consider the functional traits of the dominant plant communities in their field sites to determine how best to alleviate negative impacts on biological structure.

Researcher direct and indirect effects had stronger, negative effects on plant communities and sediment conditions in our constructed marshes (i.e., AL-CON and CA-CON) than our natural marshes (i.e., AL-NAT and CA-NAT; Figure 1). This difference is likely due to the fact that early-successional ecosystems, including constructed and restored tidal marshes, have less developed ecological memory [i.e., the information (e.g., functional traits, historical disturbance regimes) and material (e.g., seeds/rhizomes, nutrient pools) legacies of an ecosystem that impact its capacity to respond to disturbance] than natural ecosystems (Johnstone et al. 2016). For example, it is well-documented that AL-CON has less belowground plant biomass and smaller nutrient pools than AL-NAT (Tatariw et al. 2021), suggesting that the ecological memory in AL-CON limits its capacity to recover following disturbance. Similarly, we only observed subordinate species (e.g., *D. spicata* in AL and *S. bigelovii* in CA) colonizing researcher impacted plots (i.e., High Crab, Low Crab, Trampled) in natural tidal marshes, suggesting that the rhizome/seed banks in the constructed marshes are limited and reduced the capacity of these marshes’ plant communities to respond to disturbance.

Ecological memory, like other ecosystem attributes, develops through time; thus, older constructed and restored ecosystems should have greater resistance and resilience to disturbance than younger constructed and restored ecosystems (Johnstone et al. 2016). Our results support this assumption; researcher effects were less severe in the Alabama constructed marsh, which is twice as old as the California constructed marsh (Figure 1; CA-CON = 17 years, AL-CON = 34; Vittor et al. 1987; Beheshti et al. 2022). However, even after three decades, AL-CON has not developed the resiliency necessary to recover from researcher direct and indirect effects, which are still evident >2 years after the initial disturbance (Rinehart and Dybiec *personal observations*). Consequently, while research in constructed and restored ecosystems is clearly necessary to overcome historical and contemporary rates of ecosystem loss (Guan et al. 2019; Aronson et al. 2020), we advocate that researchers working in these early-successional ecosystems implement strategies to mitigate their scientific footprint (see Figure 6).

**Figure 6.**
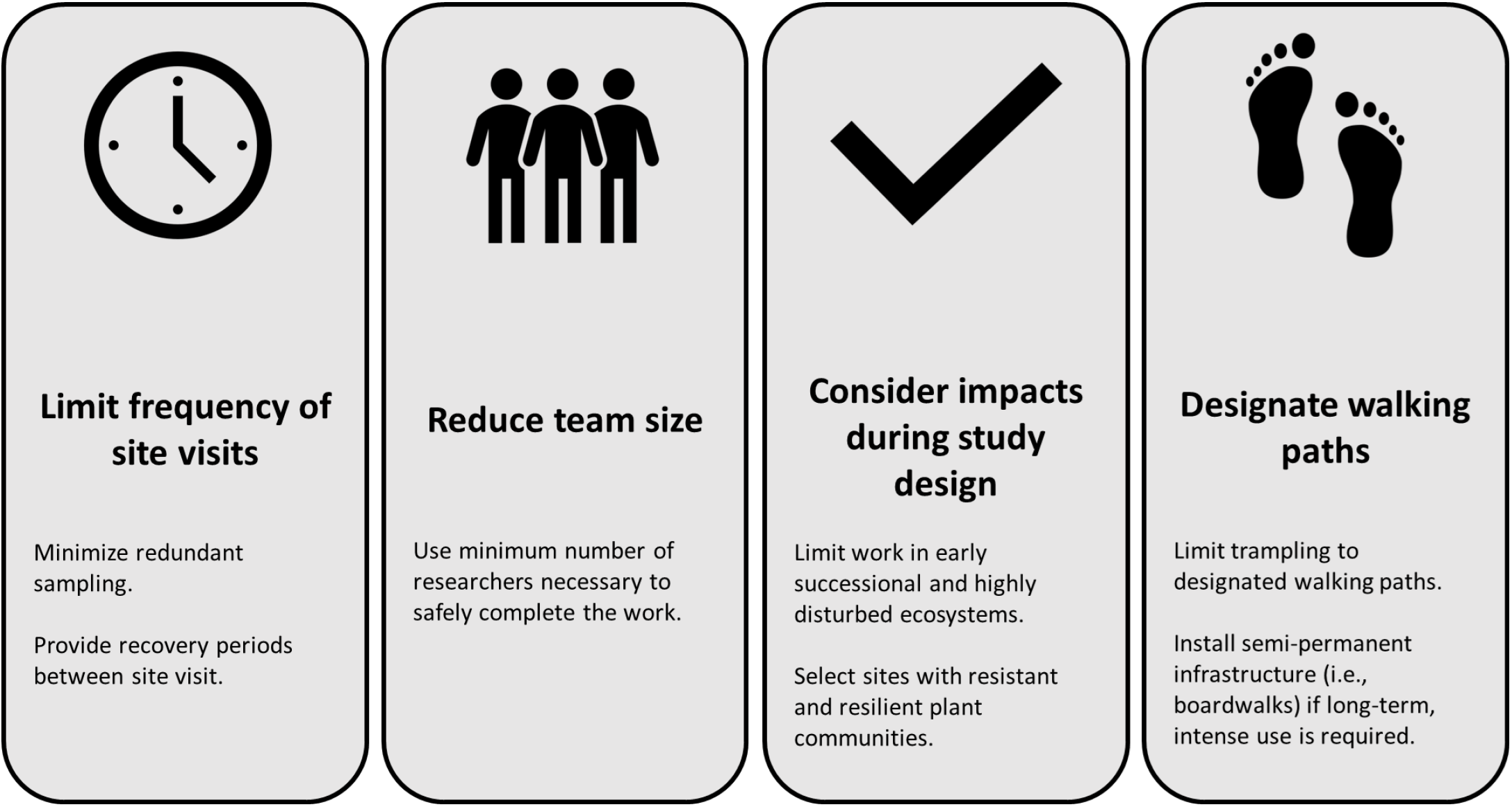
Schematic showing the four major strategies that researchers can use to minimize their scientific footprint, including limiting the frequency of site visits, reducing team size, considering impacts during experimental design, and designating researcher walking trails.

### Implications for conservation and management

It is well-known that human activities, like trampling, can affect the productivity and overall health of tidal marsh ecosystems and other critical ecosystems worldwide (Cole 1987, 1995; Goldman Martone & Wasson 2008; Pescott & Stewart 2014). However, research activities (e.g., basic research, bioassessment, etc.) are vital to understanding, managing, and conserving ecosystems, especially considering the projected impacts of climate change. Taking members of the public into these ecosystems on tours also increases support for conservation and decreases hostility between natural reserves and their residential neighbors (Davenport et al. 2007). Thus, there is a continued and paradoxical need to disturb critical ecosystems to ensure their longevity and health.

Given this, field researchers need to act as environmental stewards and work to actively reduce their personal impacts on the ecosystems they study. Past efforts have proposed several strategies for minimizing researcher impacts in specific ecosystems, like wetlands, such as practicing ‘leave no trace’, back filling soil pits, and using biodegradable materials to mark sampling locations (see Bryzek et al. 2022). Based on our study, we support these practices and seek to further promote four key strategies that field researchers can implement to reduce their scientific footprints and limit their impacts on natural and constructed systems (Figure 6).

First, we advocate that researchers work to limit the number of site visits involved in each project. For example, Walker et al. (2021a, 2021b) monitored crab burrow density and plant communities in their crab manipulation study monthly from ∼April-October in 2016-2018. However, the main data used in the final manuscripts were from the final sampling timepoint (October) of each year. Thus, these same data could have been collected with at least 21 fewer site visits (42-84 fewer trail passes), which would have reduced the scientific footprint of these field studies considerably. Similarly, in our current studies we could have further reduced our site visitations and still produced a scientifically rigorous and informative dataset. This modification would have further limited our trampling effects on plant communities, especially early in the growing season when plants are smaller and likely more vulnerable. Second, we advocate that researchers limit the size of their field teams to the minimum number of researchers needed to safely complete the research. Reducing the number of researchers in the field will limit the amount of trampling activity and further reduce other indirect researcher effects, like trash and litter (see Bryzek et al. 2022), not assessed in our current study.

Third, we strongly encourage researchers to think about the environmental impacts of their studies during the design phase. This approach will provide the ideal opportunity to reflect on which sampling timepoints and variables are essential, which sampling timepoints and variables could be eliminated, and how many researchers will be needed to complete each task in the field. Additionally, by considering environmental impacts during the design phase, researchers can think about which field sites may be the most resilient to their activities and minimize their use of sensitive ecosystems, like restored and constructed habitats, whenever possible. For instance, we may consider limiting our use of these sensitive ecosystems to only research activities that are vital to the management and conservation of the site (e.g., monitoring) and the species that inhabit it (e.g., threatened and endangered species).

Fourth, given that even low intensity trampling can damage some plant communities (Pescott & Stewart 2014), researchers should try to designate walking paths to minimize the area being disturbed by trampling (Bryzek et al. 2022). While using a single path will have dramatic negative effects on the habitat along that path, localizing those impacts to a single path, rather than multiple paths, should limit the overall impact that research teams have on their field sites. Additionally, in highly sensitive ecosystems, it may be valuable to install semi-permanent infrastructure, like boardwalks, if the research will require >200 passes of a single path, since several plant communities are reduced to <90% total cover after this level of disturbance (Cole 1987).

Our research revealed direct and indirect researcher effects to natural and constructed tidal marsh ecosystems that altered biological structure and sediment conditions for at least a year post-impact. These sorts of research impacts are likely common in other habitat types and may be more severe in early successional or restored ecosystems. However, these types of field studies are essential to understand ecological processes. Thus, our intention with this study is not to reduce the amount of field research occurring or diminish the importance of field research; rather we seek to encourage our community of field researchers to 1) think more critically about the environmental impacts of their research and 2) start implementing simple strategies (see Figure 6) that will help reduce our collective scientific footprint.

## Acknowledgements

J.D. Long provided equipment. M. Sharbaugh, A. Pasierbowicz, H. Rutter, and A. Wiggins helped collect data in the lab and field. S.C. France provided insightful discussion and resources. This work was performed (in part) at the University of California Natural Reserve System (Kendall-Frost Mission Bay Marsh Reserve), https://doi.org/10.21973/N3008B. Thanks to I. Kay for access to Kendall-Frost Marsh Reserve. G. Crozier, G. McKean, B. Vittor, S. Schroeter assisted with field site access at Fowl River and San Dieguito Lagoon. Funding was provided by the University of Alabama’s Department of Biological Sciences. This is contribution #XX of San Diego State University’s Coastal and Marine Institute Laboratory.

## Author Contributions

Conception: all authors; Design: all authors; Data collection, visualization, and interpretation: PR, SR, JMD, JW, JAC; Supervision, SR, JBW, JAC; Writing-Original Draft, SR, JW, PR, JMD; Writing-Review & Editing, All authors.

## Conflict of Interest Statement

The authors claim no conflict of interest.

## Notes

### Competing Interest Statement

The authors have declared no competing interest.

