## Appendices for "Researcher effects on the biological structure and edaphic conditions of field sites and implications for management"

* Authors contributed equally

**Appendix S1:** Initial burrowing crab caging studies.

***Alabama:***

In May 2021, we (team of 2-3 researchers) installed 10 (0.5 x 0.5 x 1 m; l x w x h m) polyethylene mesh (3.1 mm openings) cages at AL-CON and AL-NAT. All cages were placed at least 1 m apart in the needlerush (*Juncus roemerianus*) zones in areas where crab burrowing was present. Cages at AL-CON were ~5-10 m from the main channel edge. In AL-NAT, cages were ~ 30 m from the edge of a tidal creek.

We inserted cages 15 cm into the sediment, which required that we sever plant rhizomes with sharp shooters. We also adhered aluminum flashing to the top edge of each cage (flashing did not affect cage heights) and installed two pitfall traps [diameter (Ø) = 7 cm, height = 20 cm) in opposite, diagonal corners of all cages. Once installed, we randomly allocated the cages to High Crab and Low Crab treatments (n = 5 $treatment^{-1}$) and closed the pitfall traps in High Crab plots (using window screening with 1mm openings) to prevent crab removal in these plots, while standardizing the disturbance. We also inserted two decomposition bags (tea bags) and added two feldspar marker horizons (0.1 x 0.1 m, l x w) in each cage.

We visited both sites monthly from May through December 2021. During these visits, teams of 2-3 researchers repaired cages, removed burrowing crabs from Low Crab cages, and added burrowing crabs to High Crab cages. We also collected data on crab burrows (density, Ø) and plant communities (percent cover, plant heights, stem densities). In December 2021, we (team of 6 researchers) harvested the aboveground biomass from each cage (at the sediment-air interface), extracted a sediment core (Ø = 7.9 cm, depth = 10 cm) to harvest belowground biomass, extracted three sediment cores (Ø = 5 cm, depth = 10 cm) to collect samples for bulk density and sediment organic matter content, and extracted four sediment cores (Ø = 2 cm, depth

= 10 cm) to sample accretion on marker horizons. After sample collection, we removed the plastic mesh cages, but left the cage footprints marked with wooden stakes.

***Southern California:***

In May 2016, we (team of 5-6 researchers) installed 10 (0.7 x 0.7 x 0.6 m, l x w x h) plastic mesh cages (0.6 cm openings) at CA-CON and CA-NAT. All cages were placed at least 1 m apart at the ecotone between Pacific cordgrass (*Spartina foliosa*) and pickleweed (*Sarcocornia pacifica*) in areas where crab burrowing was present. In CA-CON, cages were ~10 m (horizontal distance) from the main channel's mean low low water mark. In CA-NAT, cages were ~2-3 m (horizontal distance) from the edge of a tidal creek.

We inserted cages 30 cm into the sediment, which required that we sever plant rhizomes with sharp shooters. We also adhered aluminum flashing to the top edge of each cage (flashing did not affect cage heights) and installed two pitfall traps (Ø = 7 cm, height = 20 cm) in opposite, diagonal corners of all cages. Once installed, we randomly allocated the cages to High Crab and Low Crab treatments (n = 5 $treatment^{-1}$) and closed the pitfall traps in High Crab plots (using lids) to prevent crab removal in these plots, while standardizing the disturbance.

We visited both sites every 2-3 weeks from May to October 2016. We kept the cages installed through the winter and visited them every 2-3 weeks during the 2017 and 2018 growing seasons (April-October). During these visits, teams of 2-4 researchers repaired cages, removed burrowing crabs from Low Crab cages, and added burrowing crabs to High Crab cages. We also collected data on crab burrows (density, Ø) and plant communities (percent cover, plant heights, stem densities) every three months each year of the study (2016-2018). In October 2018, we (team of 6-8 researchers) harvested the aboveground biomass from each cage (at the sediment-air interface) and extracted a 27-cm-deep sediment core (Volume ~3,980 $cm^3$) from the center of each cage to harvest belowground plant biomass. After sample collection, we removed the plastic mesh cages, but left the cage footprints marked with wooden stakes. A full description of the manipulation is detailed in Walker et al. (2021a, 2021b).

**Appendix S2:** ANOVA output tables.

**Table S1** Two-factor ANOVA outputs for the effects of researchers on plant cover and traits of the dominant plant species in Alabama (AL) and California (CA) tidal marshes.

| Region | Indicator | Site | | | Treatment | | | Site x Treatment | | |
|---|---|---|---|---|---|---|---|---|---|---|
| | | F | df | p | F | df | p | F | df | P |
| | **Plant cover** | | | | | | | | | |
| AL | Total cover | 0.0006 | 1 | 0.981 | 118.59 | 3 | **< 0.001** | 4.44 | 3 | **0.010** |
| CA | Total cover | 248.29 | 1 | **< 0.001** | 25.08 | 3 | **< 0.001** | 25.44 | 3 | **< 0.001** |
| | **Stem Density** | | | | | | | | | |
| AL | Needlerush | 0.002 | 1 | 0.964 | 25.93 | 3 | **< 0.001** | 5.22 | 3 | **0.005** |
| CA | Pacific cordgrass | 3.42 | 1 | 0.070 | 0.75 | 3 | 0.530 | 3.23 | 3 | **0.040** |
| | **Stem Height** | | | | | | | | | |
| AL | Needlerush | 0.25 | 1 | 0.620 | 24.16 | 3 | **< 0.001** | 0.29 | 3 | 0.840 |
| CA | Pacific cordgrass | 1.6 | 1 | 0.220 | 6.63 | 3 | **0.001** | 1.4 | 3 | 0.260 |

P-values in bold are significant at $\alpha = 0.05$.

**Table S2** Two-factor ANOVA output for researcher effects on belowground biomass, sediment bulk density, and sediment organic matter content in natural and constructed Alabama tidal marshes (AL-NAT and AL-CON, respectively).

| | Depth | | | Treatment | | | Depth x Treatment | | |
|---|---|---|---|---|---|---|---|---|---|
| | F | df | P | F | df | p | F | df | p |
| **Belowground biomass** | | | | | | | | | |
| AL-NAT | 3.76 | 1 | 0.061 | 2.60 | 3 | 0.069 | 2.46 | 3 | 0.080 |
| AL-CON | 1.79 | 1 | 0.190 | 2.53 | 3 | 0.075 | 3.25 | 3 | **0.034** |
| **Bulk density** | | | | | | | | | |
| AL-NAT | 7.748 | 3 | **<0.001** | 0.749 | 3 | 0.527 | 1.032 | 9 | 0.425 |
| AL-CON | 3.603 | 3 | **0.018** | 4.662 | 3 | **0.005** | 0.360 | 9 | 0.950 |
| **Organic matter** | | | | | | | | | |
| AL-NAT | 9.48 | 3 | **<0.001** | 1.11 | 3 | 0.350 | 1.10 | 9 | 0.376 |
| AL-CON | 0.741 | 3 | 0.531 | 11.404 | 3 | **<0.001** | 0.163 | 9 | 0.997 |

P-values in bold are significant at α =0.05.

**Appendix S3:** Supplemental figures

**Figure S1:**

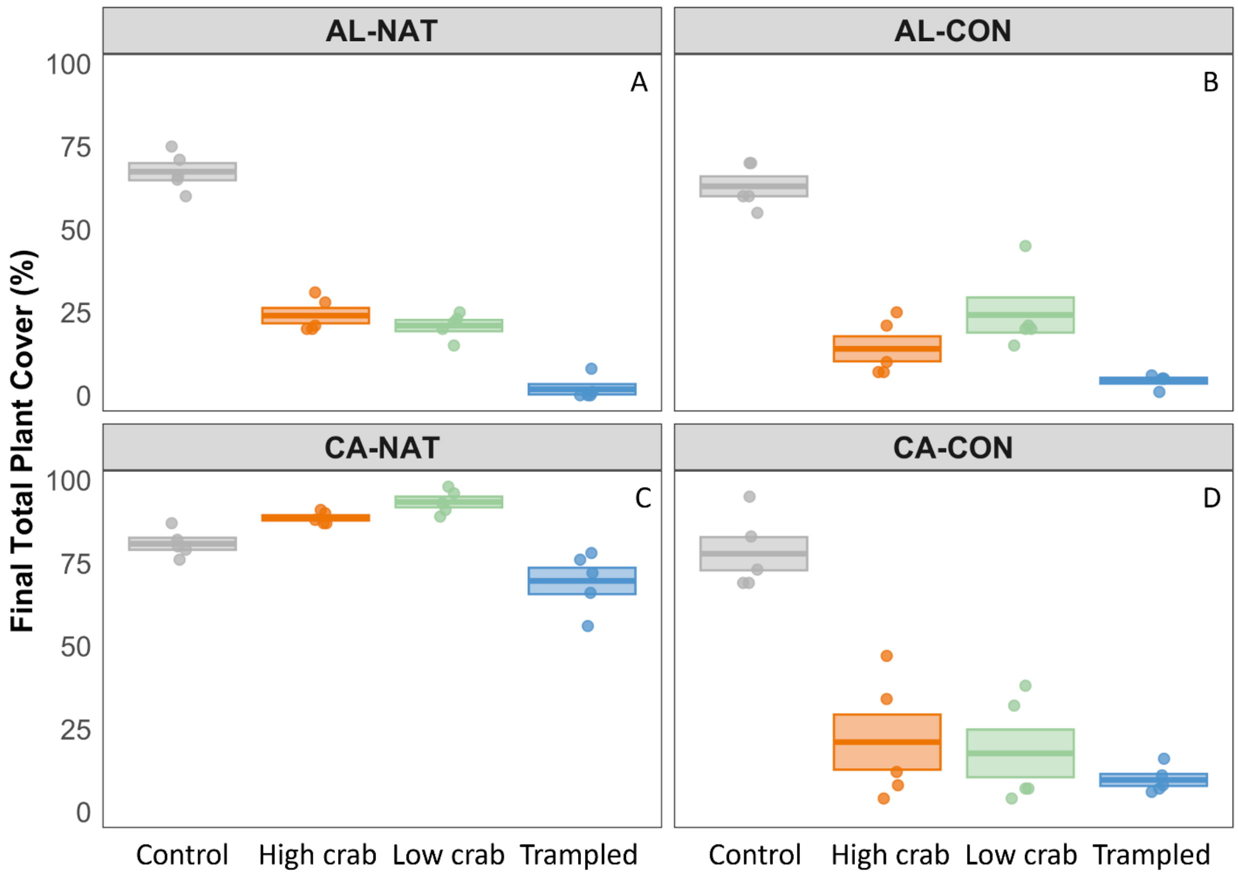


**Figure S1** Total plant percent (%) cover at the end of the growing season in all treatments (Control, High Crab, Low Crab, and Trampled) in the A) Alabama natural (NAT) marsh, B) Alabama constructed (CON) marsh, C) California NAT marsh, and D) California CON marsh. Lines inside boxes are mean values, box limits represent ± 1 SE. Points represent raw data.

**Figure S2:**

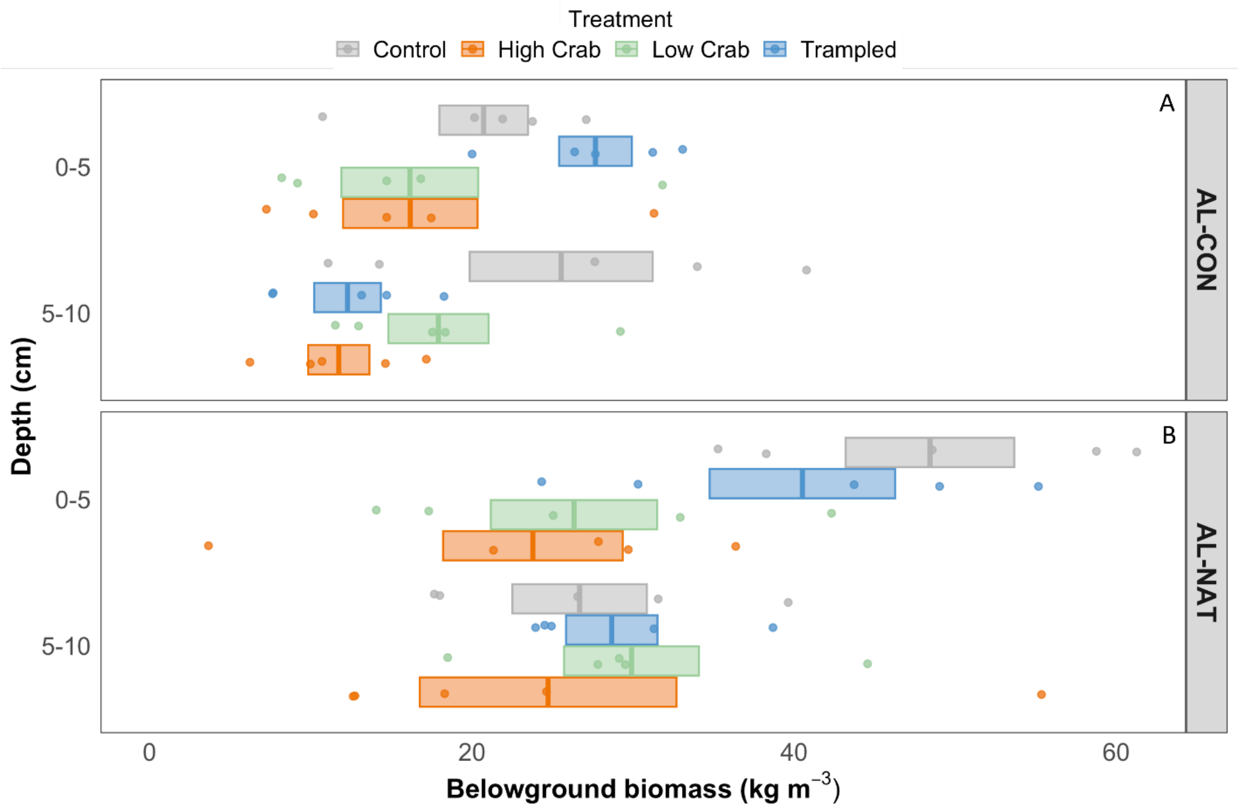


**Figure S2** Belowground biomass (kg $m^{-3}$) across depths in all treatments (Control, High Crab, Low Crab, and Trampled) at the A) Alabama constructed tidal marsh (AL-CON) and B) Alabama natural tidal marsh (AL-NAT). Lines inside boxes are mean values, box limits represent ± 1 SE. Points represent raw data. There were no significant ($\alpha = 0.05$) outputs for Tukey HSD tests at each marsh.

**Figure S3:**

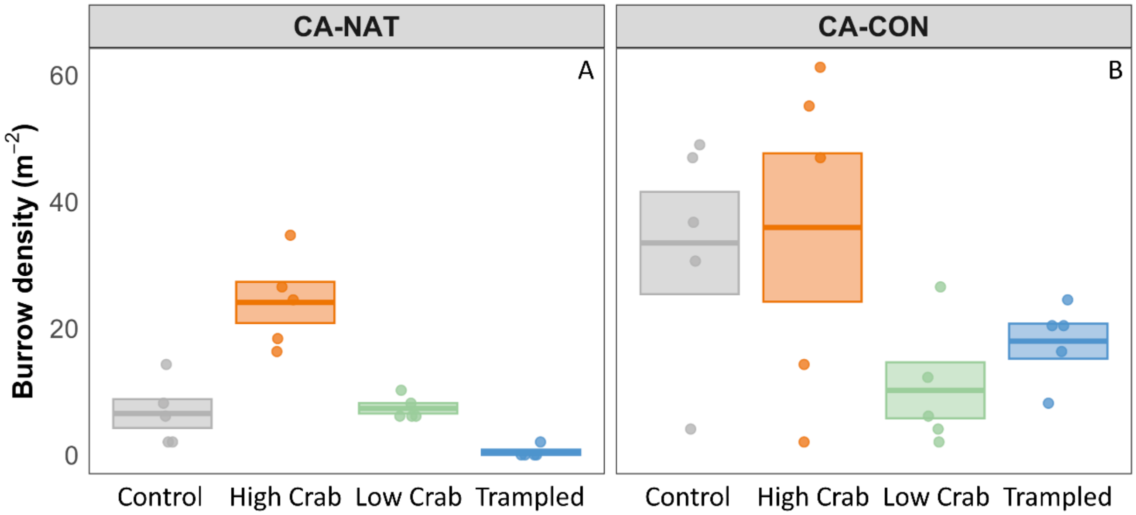


**Figure S3** Crab burrow density (m$^{-2}$) at the end of the growing season in all treatments (Control, High Crab, Low Crab, and Trampled) in the California A) natural (NAT) marsh and B) constructed (CON) marsh. Lines inside boxes are mean values, box limits represent ± 1 SE. Points represent raw data.

**Figure S4:**

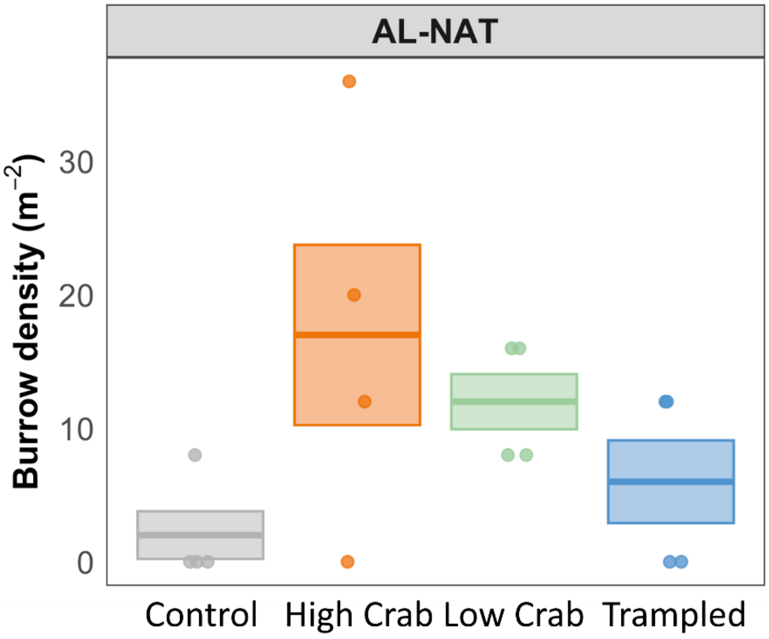


**Figure S4** Crab burrow density (m$^{-2}$) at the end of the growing season in all treatments (Control, High Crab, Low Crab, and Trampled) in the Alabama natural (NAT) marsh. Lines inside boxes are mean values, box limits represent ± 1 SE. Points represent raw data.
